# Automated osteosclerosis grading of clinical biopsies using infrared spectroscopic imaging

**DOI:** 10.1101/692434

**Authors:** Rupali Mankar, Carlos E. Bueso-Ramos, C. Cameron Yin, Juliana E. Hidalgo-Lopez, Sebastian Berisha, Mustafa Kansiz, David Mayerich

**Affiliations:** Department of Electrical and Computer Engineering, University of Houston; Department of Hematopathology, MD Anderson Cancer Center; Photothermal Spectroscopy Corp., Santa Barbara, CA

## Abstract

Osteosclerosis and myefibrosis are complications of myeloproliferative neoplasms. These disorders result in excess growth of trabecular bone and collagen fibers that replace hematopoietic cells, resulting in abnormal bone marrow function. Treatments using imatinib and JAK2 pathway inhibitors can be effective on osteosclerosis and fibrosis, therefore accurate grading is critical for tracking treatment effectiveness. Current grading standards use a four-class system based on analysis of biopsies stained with three histological stains: hematoxylin and eosin (H&E), Masson’s trichrome, and reticulin. However, conventional grading can be subjective and imprecise, impacting the effectiveness of treatment. In this paper, we demonstrate that mid-infrared spectroscopic imaging may serve as a quantitative diagnostic tool for quantitatively tracking disease progression and response to treatment. The proposed approach is label-free and provides automated quantitative analysis of osteosclerosis and collagen fibrosis.

## Introduction

Myeloproliferative neoplasms (MPN) are a group of heterogeneous hematologic malignancies affecting the proliferation and expansion of one or more hematopoietic lineages. This dysregulation is thought to be a consequence of genetic abnormalities at the level of stem/progenitor cells. Myelofibrosis is an increase in the amount and density of extracellular matrix proteins that provide the structural network upon which hematopoiesis occurs. This increase can vary from a focal, loose network of reticulin fibers to diffuse, dense, and markedly thickened fibers associated with collagen fibrosis and osteosclerosis. Accurate grading of myelofibrosis (MF) and osteosclerosis is an important component of assessing disease staging and prognosis. Grading is traditionally performed by pathologists based on a four-grade European Myelofibrosis Network (EUMNET) scoring system.^1^ EUMNET scoring uses bone marrow (BM) biopsies stained with H&E, reticulin, and trichrome. In last WHO classification, EUMNET classification was described with collagen and osteosclerosis in separate scale for grading.^2^ However, traditional diagnosis is expensive, difficult to quantify, and imprecise due to inter-observer variations and lack of standard assessment methods, making it challenging to track successful treatment.^3^ Recent advances in virtual microscopy enable pathologists to digitize high resolution histological images, which has encouraged the development of computer-assisted tools for pathological analysis.^4^ Digital analysis increases repeatability and throughput by reducing human errors and providing deterministic results.^5–7^ Digital quantification of trabecular bone area has been proposed^8^ to evaluate the effects of cancer treatments such as imatinib mesylate.^9^ Digital analysis of stained tissues may provide more quantitative results, but is still dependent on tissue staining and prone to variations in protocols and image quality. We propose using vibrational spectroscopic imaging of unstained tissue sections, leveraging a quantitative measure of tissue molecular composition as a contrast mechanism. Infrared spectroscopic imaging, such as Fourier transform infrared (FTIR) and discrete frequency infrared (DFIR) spectroscopy is quantitative, label-free, and non-destructive. This enables both objective digital analysis and automated evaluation of histological structures for grading while preserving biopsies for downstream analysis.

Infrared (IR) absorbance is often used to identify molecular signatures in organic materials. Fourier transform infrared (FTIR) has been applied for label free characterization and classification in histopathological studies^10–16^ and has the potential to automate examinations to both save time and reduce diagnostic errors. This paper focuses on evaluating the potential for absorbance spectroscopy to automate osteosclerosis and collagen grading, particularly trabecular bone area (TBA), by defining a clinically-viable protocol for image acquisition and analysis. FTIR spectroscope has been successful at differentiating collagen sub-types (I, III, IV, V, and VI),^17^ however it has not been validated on clinical samples. Comprehensive analysis of abnormal growth of type I and type III collagen is critical for MPN prognosis, therefore we also explore pixel-level classification of these collagen sub-types using clinical biopsies. Finally, we develop a set of metrics that provide promising results for automated grading of both osteosclerosis and collagen EMN components.

## Background

Bone marrow (BM) is a soft spongy tissue located within bone cavities and is primarily responsible for new blood cell formation (hematopoiesis). Classification of MPNs is based on clinical, laboratory, morphologic, and genetic findings at the time of initial presentation, prior to any definitive therapy for the myeloid neoplasm. Diagnosis of hematologic disorders relies on characterization of hematopoietic cells and collagen content within the marrow - specifically the size and shape of bone trabeculae, as well as the fiber content of type I collagen (based on trichrome staining) and type III collagen (based on reticulin staining). We limit our focus to osteosclerosis and collagen type I grading, since type III collagen fibers require high-resolution instrumentation that is only recently commercially available in the form of Optical Photothermal Infrared (O-PTIR) imaging.^18–20^

### Osteosclerosis Grading

Osteosclerosis is a disorder characterized by abnormal trabecular bone growth often accompanied by fibrosis.^21^ The semiquantitative grading of osteosclerosis is shown in Figure 1. The scoring for osteosclerosis grading according to the World Health Organization (WHO) is as follows: Grade 0, regular bone tuberculae (distinct paratrabecular borders); Grade 1, focal budding, hooks, spikes, or paratrabecular apposition of new bone; Grade 2, diffuse paratrabecular formation of new bone with thickening of trabeculae, occasionally with focal interconnections; Grade 3, extensive interconnecting meshwork of new bone with overall effacement of marrow spaces.^1,2,22^ Accurate grading of osteosclerosis and fibrosis is important for precise staging and prognosis of MPN.^22,23^ In patients with chronic myelogenous leukemia, precise evaluation of trabecular bone area is important for assessing treatment progress and resistance.^8^ Objective classification can aid characterization of MPNs,^3^ since pathological analysis of TBA is prone to inter-observer variability. Digital analysis using H&E-stained tissue sections has been proposed to address this problem^3,8^ by manually outlining trabecula to estimate TBA with respect to the total biopsy area. These studies show increased throughput and precision in TBA measurement. However, manual outlining is laborious and time-consuming. Automating precise segmentation of trabecular bone from H&E-stained BM biopsies is complicated by low staining quality and variability in staining.^24^

**Figure 1:**
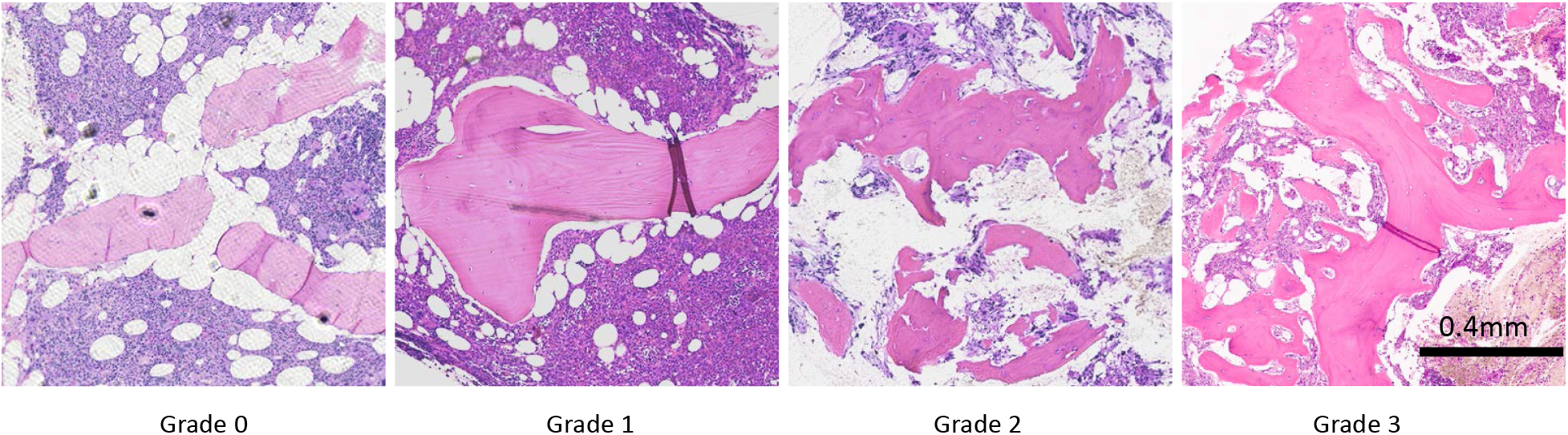
Osteosclerosis grading in bone marrow tissue. From left to right bone marrow biopsy sections for each grade from grade 0 to grade 3 and each section is stained with H&E stain and imaged at 10x magnification. Grade 0 shows normal trabecular bone in BM. Grade 1 shows initial trabecular apposition though focal budding. Grade 2 shows abnormal growth with thickening and diffused trabecular structure. Grade 3 shows extensive interconnected new bone growth with effacement of marrow space.

### Collagen Grading

The most recent WHO grading score for collagen deposition is based on a four-grade scheme using Masson’s trichrome to label type I collagen (Figure 2).^22^ Grade 0 corresponds to normal BM, with type I collagen limited to perivascular regions only. According to WHO, Grade 0 has perivascular collagen only (normal); Grade 1 shows focal paratrabecular or central collagen deposition with no connecting meshwork; Grade 2 shows Paratrabecular or central deposition of collagen with focally connecting meshwork or generalized paratrabecular apposition of collagen; and Grade 3: Diffuse (complete) connecting meshwork of collagen in > 30% of marrow spaces.^2,22^ Automating conventional histology with Masson’s trichrome staining is also challenged by staining quality (i.e. weak staining, overstaining).

**Figure 2:**
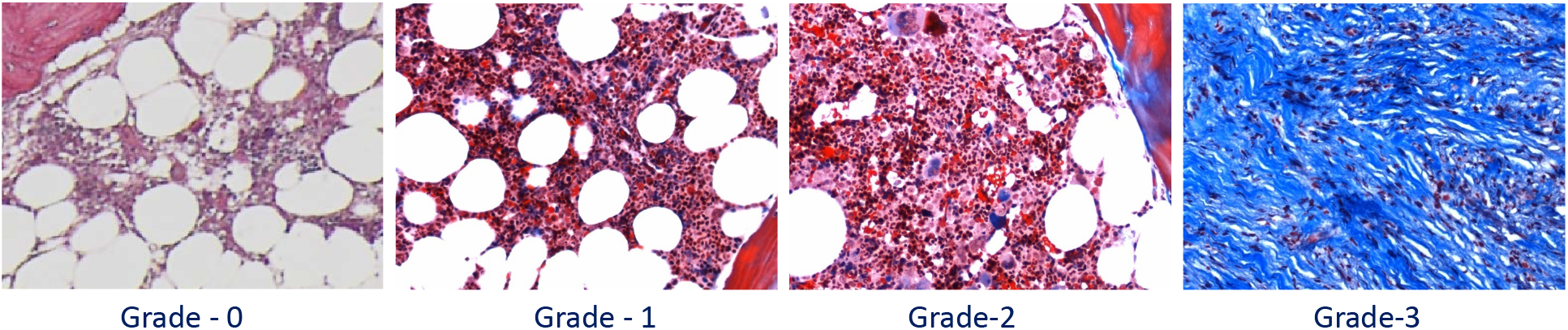
Collagen deposition grading in bone marrow tissue. From left to right bone marrow biopsy sections for each grade from grade 0 to grade 3 and each section is stained with Masson’s trichrome stain and imaged at 10x magnification. Grade 0 is normal bone marrow with only perivascular type I collagen (blue). Grade 1 shows minimal presence of type I collagen in the central are of bone marrow. Grade 2 shows paratrabecular and prominent central deposition with interconnecting collagen fibers. Grade 3 shows extensive interconnected type I collagen fibers.

## Approach

Cell and tissue classification using FTIR imaging is a mature area of research,^25–29^ however specific clinical applications are rarely seen. We propose a protocol for automated classification of trabecular bone and collagen and calculation of TBA. We then demonstrate promising and consistent results for automated osteosclerosis and collagen grading, achieving comparable accuracy to human expert evaluation based on quantitative molecular and spatial measurements.

Osteosclerosis scoring is based on evaluation of morphological variations (including size, shape, and spacing) of trabeculae across a biopsy. Accurate segmentation and quantification of trabeculae provide faster and more consistent diagnosis. We test two substrates for clinical practice: (1) standard glass slides, which are compatible with current histological practice, but opaque to IR radiation <2500 cm^−1^,^30^ and (2) IR reflective slides that provide the entire spectrum between 900 to 3900cm^−1^. Testing with both substrates is important for translational work, since the ability to grade samples using features within the glass transmission window eliminates the need for specialized substrates, such as calcium fluoride or low-emissivity glass coatings, which would require changes to existing sample preparation protocols.

Both osteosclerosis and collagen grading use a two-step classification process. Pixel-level classification is used to identify histologically relevant tissue types.^31^ For each classified biopsy, a set of spatial metrics is computed to mimic the evaluation process outlined in current pathological guidelines.^22^ A support vector machine (SVM) then maps these metrics to generate the final grading.

Three osteosclerosis metrics quantify the size and shape of trabecular cross-sections relative to the biopsy area. The first metric is TBA, which calculates the total trabecular bone cross-section relative to the biopsy area. A *diffusion metric* quantifies the scattered growth of trabeculae in higher-grade samples. An average spacing metric quantifies the regularity of interstitial spacing between trabecula. Digital scoring of type I collagen is based on its overall biopsy coverage as well as the coverage of the largest interconnected collagen cluster. This enables us to automate osteosclerosis grading and provide quantitative analysis for collagen deposition grading without any histochemical staining.

## Materials and Methods

### Sample Preparation

All tissue sections are de-calcified using 5% formic acid, fixed in 10% neutral buffered formalin, and embedded in paraffin. Sections are cut at 5 μm thickness and placed on either reflective low-emmissivity glass (Kevley Technologies) or standard glass. Anonymized tissue microarrays (TMAs) were purchased from a public repository (AMSBIO). De-identified bone marrow biopsy samples were collected from patients with MPN under an Institutional Review Board (IRB) approved protocol.

Adjacent sections were placed on standard glass for validation using standard histological stains. TMAs were assessed by an expert pathologist to annotate collagen and estimate trabecular bone area. Clinical biopsies were annotated for collagen deposition and evaluated using standard grading protocols.^22^

### Image Acquisition

FTIR images for BM TMAs were collected using a Cary 620 FTIR microscope (Agilent Technologies) in standard magnification mode using a 15× 0.65 NA objective. Spectra were acquired in the range of 900 to 3900 cm^−1^ at a spectral resolution of 8 cm^−1^ for tissues on IR reflective substrates (collected in transflectance mode) and in the range of 2800 to 3900 cm^−1^ for tissues on histology glass slides (collected in transmission mode) for osteosclerosis grading.

### Tissue Pixel-Level Classification

All mid-IR images were processed using standard protocols,^32^ which include baseline correction and normalization to the Amide I band (1650 cm^−1^) to compensate for scattering and variations in thickness and density. Pixel-level annotations were created for ground truth images based on adjacent histological sections for trabecular bone, collagen, and stroma using both H&E and Masson’s trichrome as references. A supervised feature selection algorithm was applied to identify prominent spectral differences between classes. Feature selection was performed with a previously published genetic algorithm leveraging linear discriminant analysis.^29^ A Random Forest (RF) classifier^11^ was trained on the selected features to differentiate between trabecular bone, collagen, stroma, and background pixels. The RF used 100 untruncated trees for classification.

### Osteosclerosis Grading

Trabecular bone area was calculated from classified IR images by integrating pixels identified as trabecular bone. Subcortical thickening of the trabecular bone was discarded by following guidelines for osteosclerosis grading.^22^ The area of segmented trabecular bone was calculated relative to the total tissue area to determine TBA%. Total biopsy area was estimated by integrating stromal pixels as well as adipose tissue. Since adipose tissue is removed during paraffin infiltration as part of the tissue preparation process, the resulting holes were filled and reconnected with binary morphological operations in MATLAB (*imfill* and *imclose*).

A *diffusion metric* was calculated by counting the number of contiguous trabecular bone structures covering 80% of the total TBA. Remaining pixels were ignored to avoid considering tiny trabecular structures caused by tissue preparation and classification errors. An *average spacing metric* describing regularity in trabecular bone spacing was calculated using a distance transform on segmented trabecular bone.

Grading was validated by classifying all clinical osteosclerosis samples into one of the four grades. The three metrics were computed for all clinical samples and an SVM classifier was trained and tested using leave-one-out cross-validation.

### Collagen Grading

The percentage of type I collagen fiber (CD%) was computed by integrating the total number of collagen pixels, excluding the perivascular pixels, with respect to total tissue area. The area of the largest connected collagen cluster was calculated, providing a characterization of the extensive interconnected collagen meshwork used to differentiate grades 2 and 3 according to guidelines.

## Results and Discussion

### FTIR Imaging on Glass Slides

FTIR imaging generally provides a spectral range of 800 to 4000cm^−1^^32^ which consists of a chemically rich fingerprint region (800 to 1800cm^−1^) separated from a functional group region (2550 to 3500cm^−1^). The intermediate 1800 to 2550cm^−1^ region is primarily composed of atmospheric absorbance. Conventional glass slides heavily absorb in the fingerprint region, making them generally unsuitable for FTIR spectroscopy. However, some histological studies suggest that glass is viable for some applications^30,33^ requiring only functional groups associated with stretching vibrations including CH, OH, and NH. This potentially contains absorbance peaks useful for trabecular bone and collagen characterization, and may aid clinical adoption by maintaining current sample preparation protocols.^34^

While the trabecular bone matrix contains both collagen fibers and minerals, almost 90% of the organic substance is collagen. De-calcification of BM tissue, which is a common step before paraffin embedding, removes minerals. Figure 3 shows mean spectra from both the trabecular bone and stroma classes. Collagen features are seen in trabecular bone in the fingerprint region (1200, 1240, 1282 and 1403 cm^−1^) and in the functional OH/NH range. The presence of collagen IV in trabecular bone introduces distinguishing features in the OH/NH range, including extra shoulders and a prominent shift to higher wavenumbers. Trabecular bone is also dense when compared to stroma, resulting in lower paraffin concentration, as is clearly seen in the spectrum at the CH stretching (2800 to 3000cm^−1^) and CH bending (≈1450 cm^−1^) bands. Features such as CH stretching (2800 to 3000cm^−1^) and collagen in the OH/NH provide a compelling case for studying classification on glass substrates for osteosclerosis grading.

**Figure 3:**
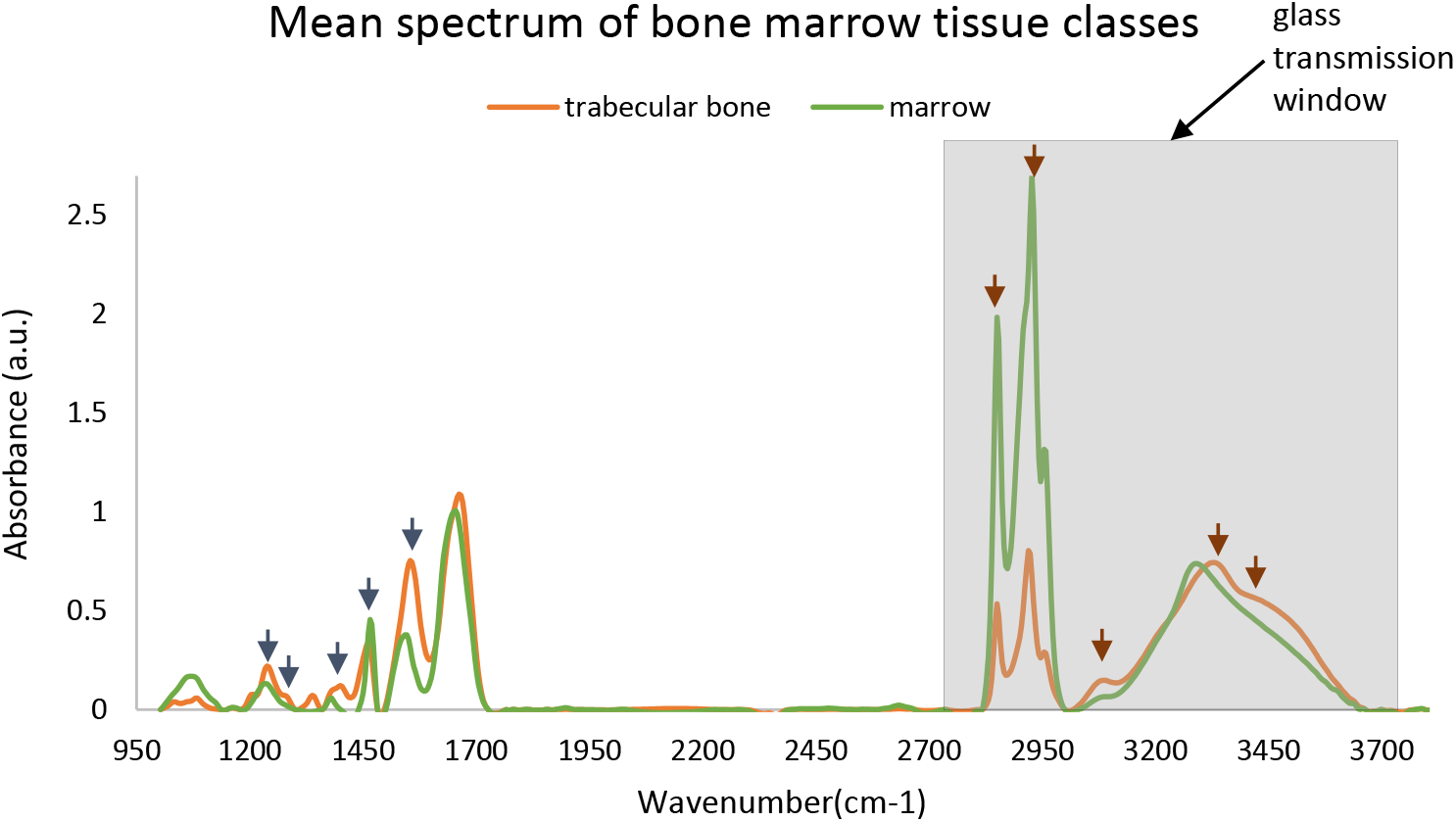
Mean spectra for two classes: trabecular bone and marrow (everything other than trabecular bone in BM) from paraffin embedded bone marrow tissue microarrays. Optimal features selected by GA-LDA^29^ for classification of trabecular bone and stroma are shown with short arrows in both fingerprint region (dark blue arrows) and in glass transmission window (brown color)

Adjacent sections of normal TMAs were imaged on both low-emmissivity glass and standard histological glass slides. Classifiers were trained independently on both data sets, producing comparable results as shown in Figure 4(b) and 5(b). Results show that classification on spectra within the glass transmission window does not affect classification results, suggesting that FTIR-based osteosclerosis grading is compatible with standard histology slides.

**Figure 4:**
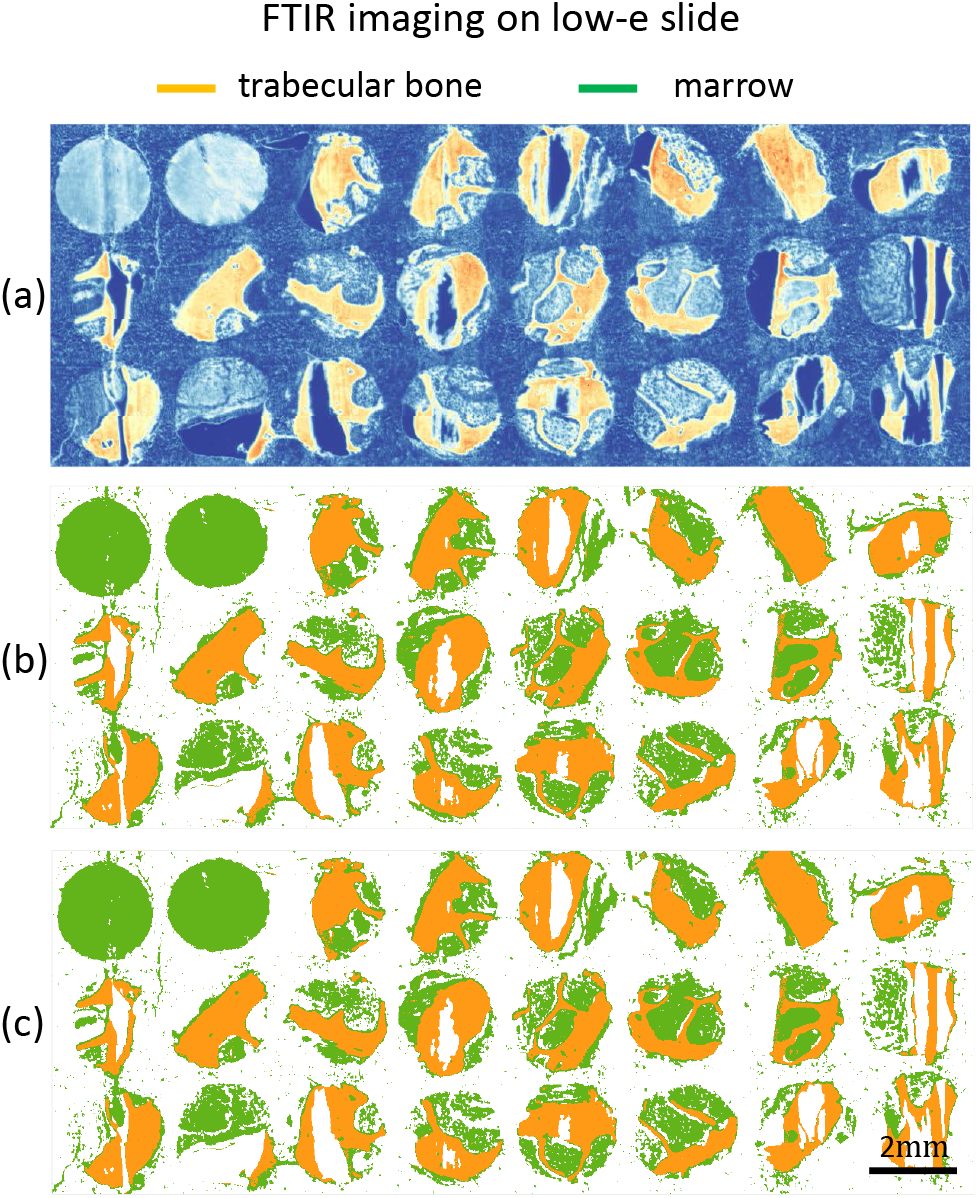
TMA of bone marrow tissue cores is imaged with FTIR on low emissivity coated glass slide. (a) Band image at 1650 cm^−1^, (b) Classification results for two classes trabecular bone and marrow (everything in BM other than trabecular bone) of bone marrow tissue core using the entire 187 features, and (c) Classification results using 5 optimal features selected using the GA-LDA algorithm.^29^

**Figure 5:**
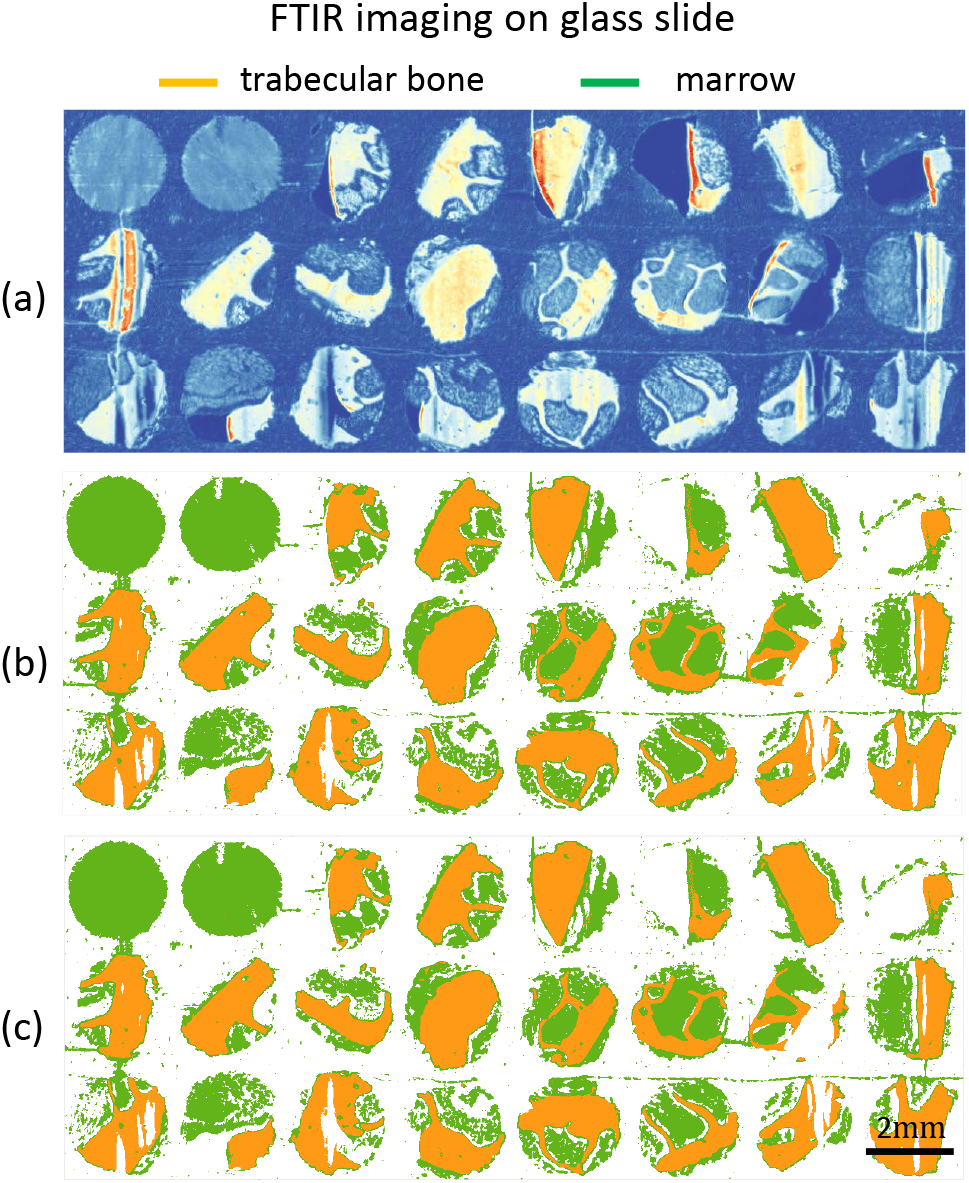
TMA of bone marrow tissue cores are imaged with FTIR glass slides. (a) FTIR band image at 3300 cm^−1^, (b) Classification results for trabecular bone and marrow (everything in BM other than trabecular bone) using all 76 spectral features in glass transmission region, and (c) Classification results using optimal 5 features selected using GA-LDA algorithm.^29^

### DFIR imaging

Our previous FTIR studies show that a minimal number of optimal features are sufficient for classification of most of the histological classes.^29^ Optimal feature selection corresponding to histological classes provides the potential for future discrete-frequency imaging methods. With prior knowledge of these features, imaging time and file size can be dramatically reduced with new laser-based imaging systems.^35^ The GA-LDA algorithm^29^ was used to select five optimal bands (Figure 3) before classification (Figure 4c and 5c). In cases of imaging on both Low-E and glass slides, comparable results were achieved using only five features, likely because spectral features between histological classes were very distinct. This suggests that a minimal number of bands from **either** the fingerprint or functional group regions are sufficient. However, currently available commercial DFIR instruments only operate in the fingerprint region, removing the possibility for discrete frequency imaging but potentially increasing imaging speed by ≈ 20×.

Optical photothermal infrared (O-PTIR) imaging provides another method for acquiring fingerprint spectra on glass substrates. While O-PTIR imaging systems are still relatively new, they provide significantly better resolution than FTIR and DFIR systems at the cost of acquisition speed.

### Osteosclerosis Grading

Our study of clinical osteosclerosis grading is retrospective. BM tissue sections were acquired from 5 patients for each grading and imaged using FTIR. Five features were selected using GA-LDA^29^ and a random forest classifier was applied to identify trabecular bone (Figures 6 and 7). Nearby sections of tissue were stained using H&E for pathological comparison. Osteosclerosis results are shown for 19 of 20 samples, as one of the samples of grade 2 exhibited fragmented trabecular bone and was removed from the study based on standard guidelines. Box plots for our three proposed metrics, along with combinations of these metrics, are shown (Figure 8). TBA% clearly separates grade 3 samples (Figure 8a), and is an important metric for assessing response to therapy for diseases such as leukemia.^8^ The diffusion metric (N) distinguishes grade 2 (Figure 8b) since the number of smaller and diffused trabecular bone structures increases from grade 0 to grade 2. In grade 3, irregularly grown trabecula fuse together to form large structures. Regular spacing between trabeculae, which is an important characteristic of grade 0 and grade 1, is quantified using the average spacing (AS) metric (Figure 8c). Higher AS values are seen for grades 0 and 1, while lower AS is seen for grades 2 and 3.

**Figure 6:**
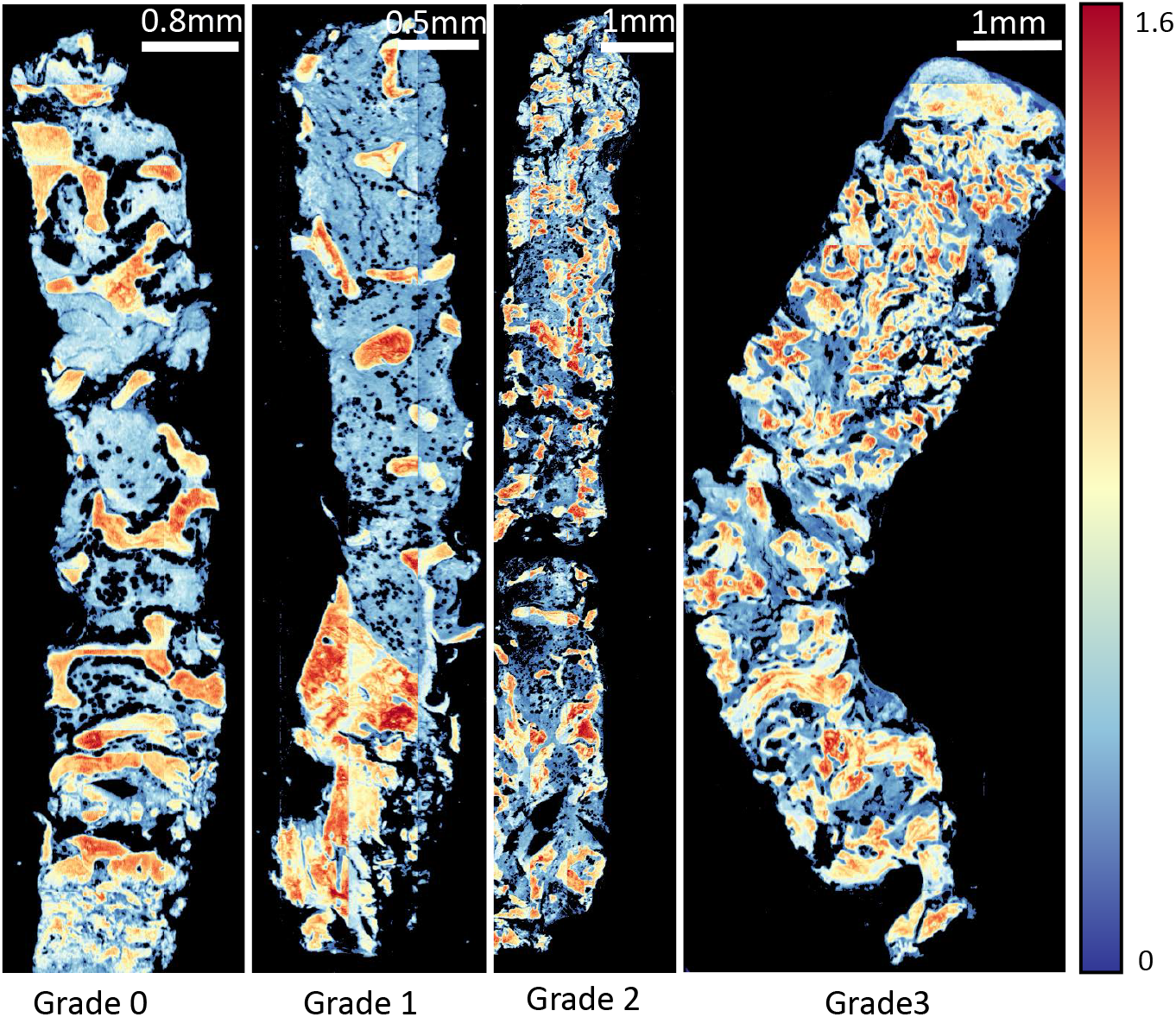
Proposed method of imaging. Trabecular bone area measurement is applied on clinical samples of bone marrow biopsy example from each osteosclerosis grade (0-3). FTIR imaged tissue at 1650 cm^−1^ wavenumber is shown for each grade.

**Figure 7:**
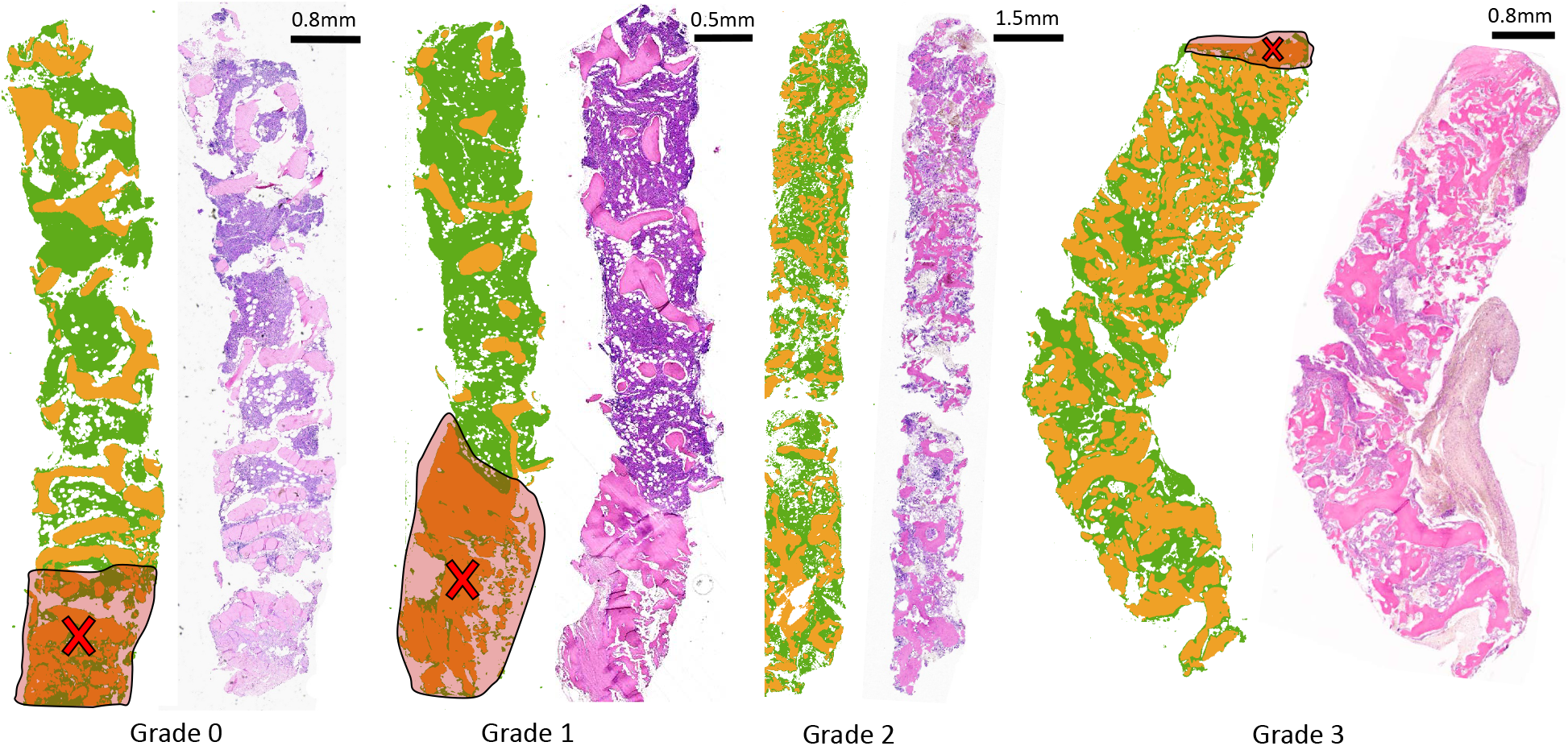
Classification result from the proposed imaging method is visually compered with traditional H&E stain. Trabecular bone area measurement is applied on clinical samples of bone marrow biopsy from each osteosclerosis grade (0-3). Results of classification (trabecular bone in orange and stroma in green) of FTIR imaged data for each grade section and corresponding H&E stained adjacent tissue section from the same tissue block (right) are shown here. Increase in trabecular bone area from grade 0 to grade 3 can be easily quantified using digitally classified images shown in the middle for each section. Cross-marks shown on the images indicate crushing artefacts on the tissue which are ignored while grading tissue sections.

**Figure 8:**
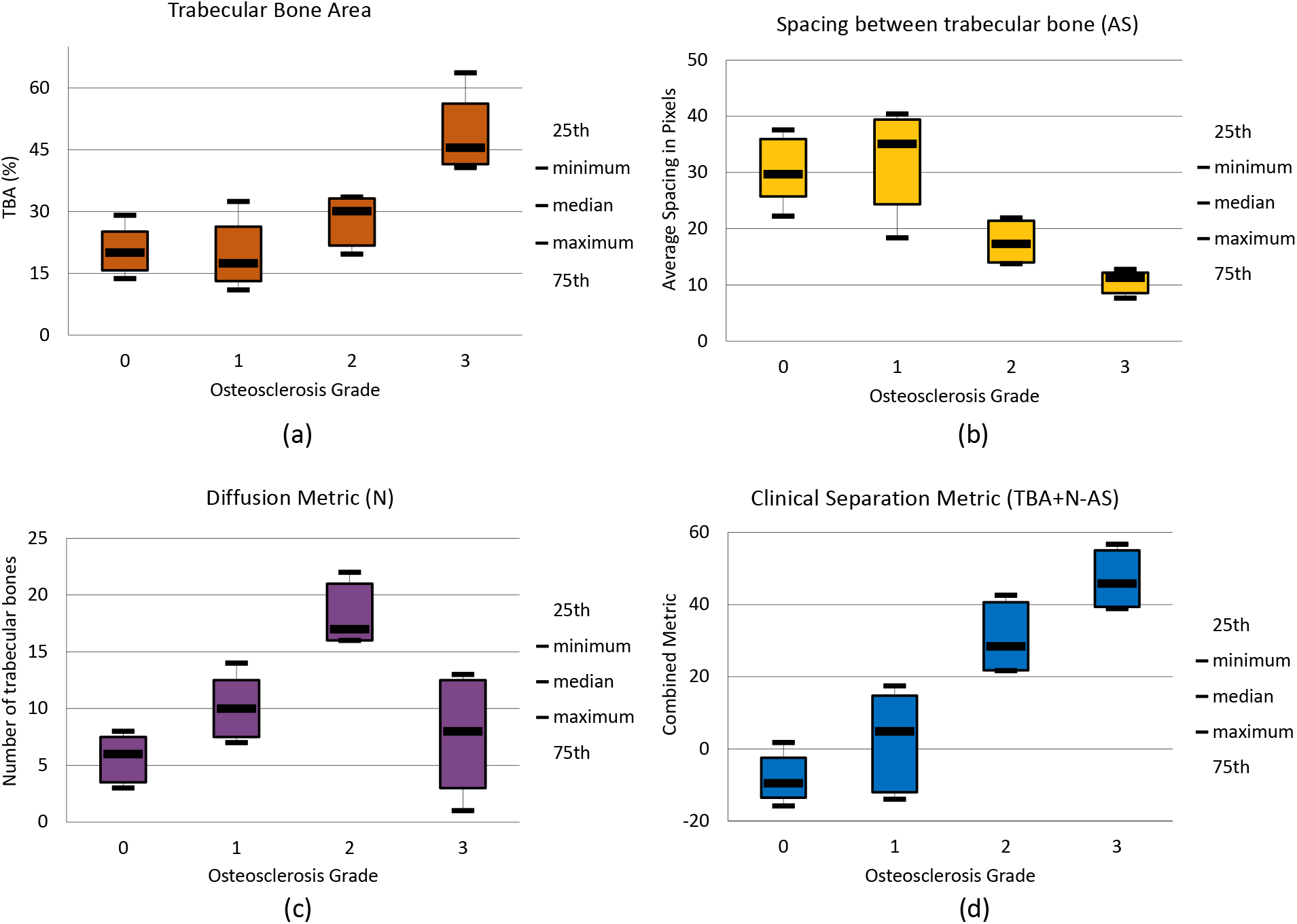
Box plots for the proposed metrics for automated osteosclerosis grading based on 19 tissue samples: (a) Trabecular bone area (TBA) - Percentage of trabecular bone area with respect to total tissue area (b) Diffusion metric (N) - count of non-fragmented trabecular bone structures contributing (c) Spacing between trabecular bone structures (AS) - Spacing is defined in pixels where pixel size is 5.5 μm^2^ (d) Clinical separation metric (CS) - combination of three metrics mentioned earlier (TBA+N-AS) clearly separates samples in two groups one with grades 0 and 1 samples and other with grades 2 and 3 samples.

Two principle features were extracted from three metrics proposed for quantification of trabecular bone using PCA and shown using scatter plots (Figure 9). Clear decision boundaries can be seen for grades 2 and 3, however separation between grades 0 and 1 is more challenging. Automated digital grading for 19 BM sections is achieved by classifying BM sections based on extracted features using SVM classifier. The grading accuracy using cross-validation on 19 BM samples is 84.4%. Given the variability of trabecular bone coverage in adjacent sections of grade 1 samples (i.e. Figure 7), it is possible that the discrepancies in grading are the result of sampling bias due to sectioning and therefore require further study.

**Figure 9:**
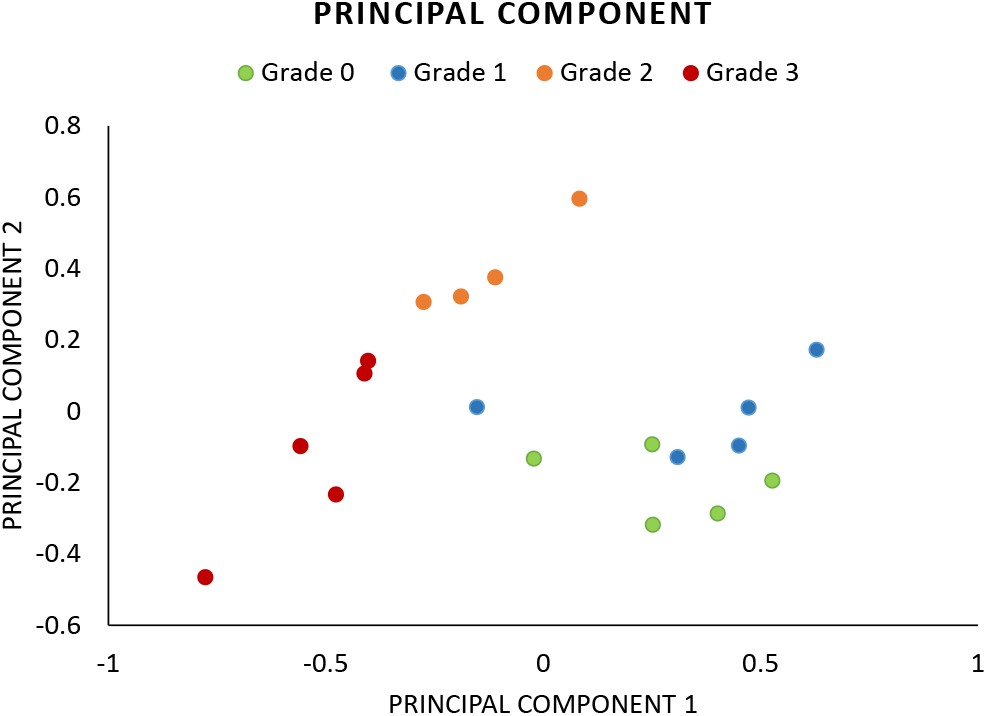
Scatter plot using first two principle component scores. Feature extraction from proposed three metrics (TBA, AS and N) gives clear separation among samples of each grades with exception of few samples from grade 0 and grade 1.

## Collagen Deposition/Collagen Fibrosis Grading

Differentiation of type I and type IV collagen (trabecular bone) requires spectral information from the fingerprint region.^17^ Conventional FTIR imaging is therefore used for collagen deposition grading. Since the spectral differences are subtle (Figure 10), BM images are collected at a spectral resolution of 4 cm^−1^ in reflection mode for collagen deposition grading.

**Figure 10:**
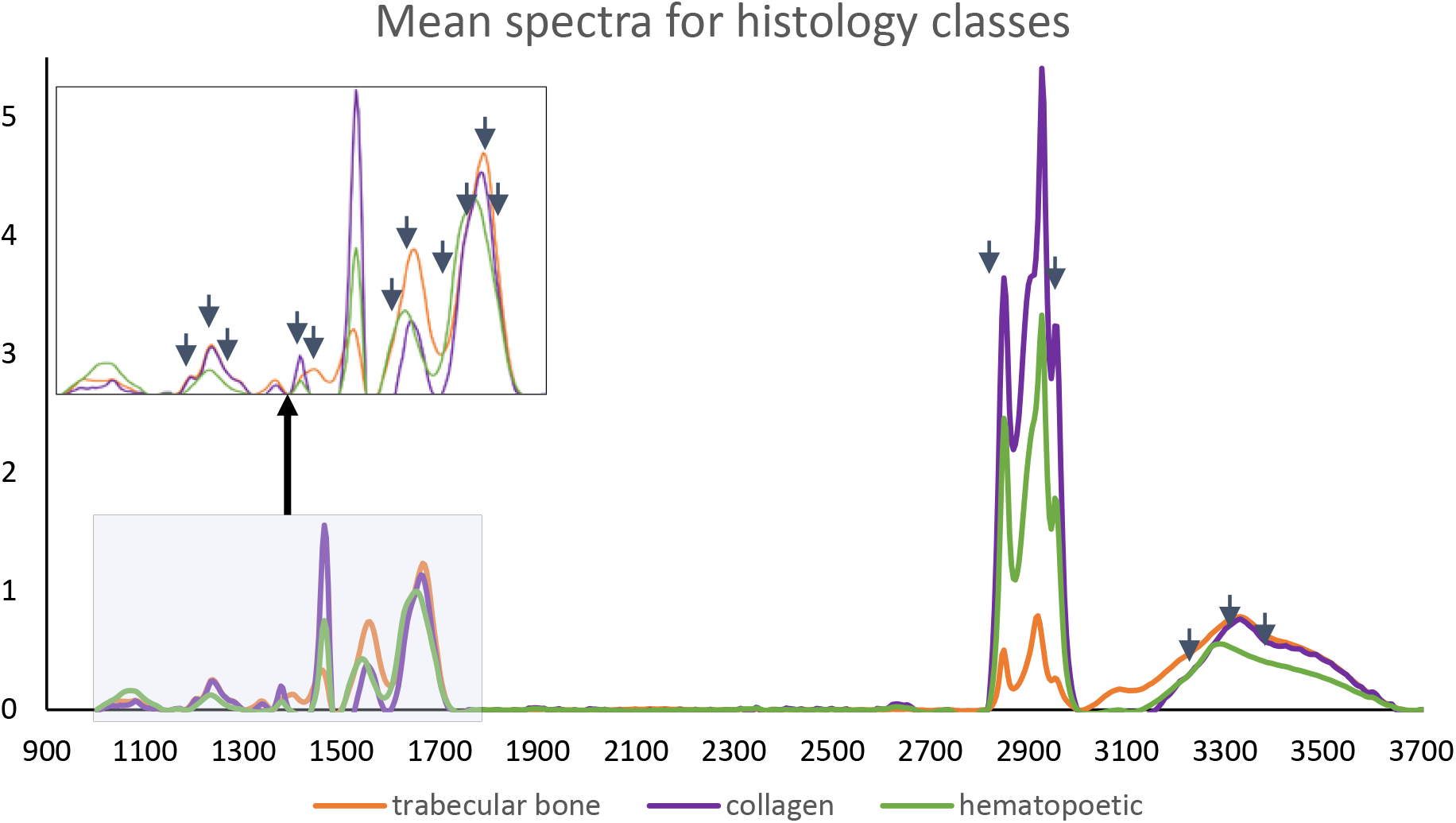
Mean absorbance IR spectra from pre-processed hyperspectral images of bone marrow biopsies for three histological classes: trabecular bone, collagen (type I) and hematopoietic cells. Arrows indicate optimal features selected using GA-LDA.

We classify using 16 optimal features to achieve an accuracy of 99.17% for three classses: type I collagen, trabecular bone, and hematopoietic cell (Figure 11). Results are comparable to adjacent sections stained with Mason’s trichrome.

**Figure 11:**
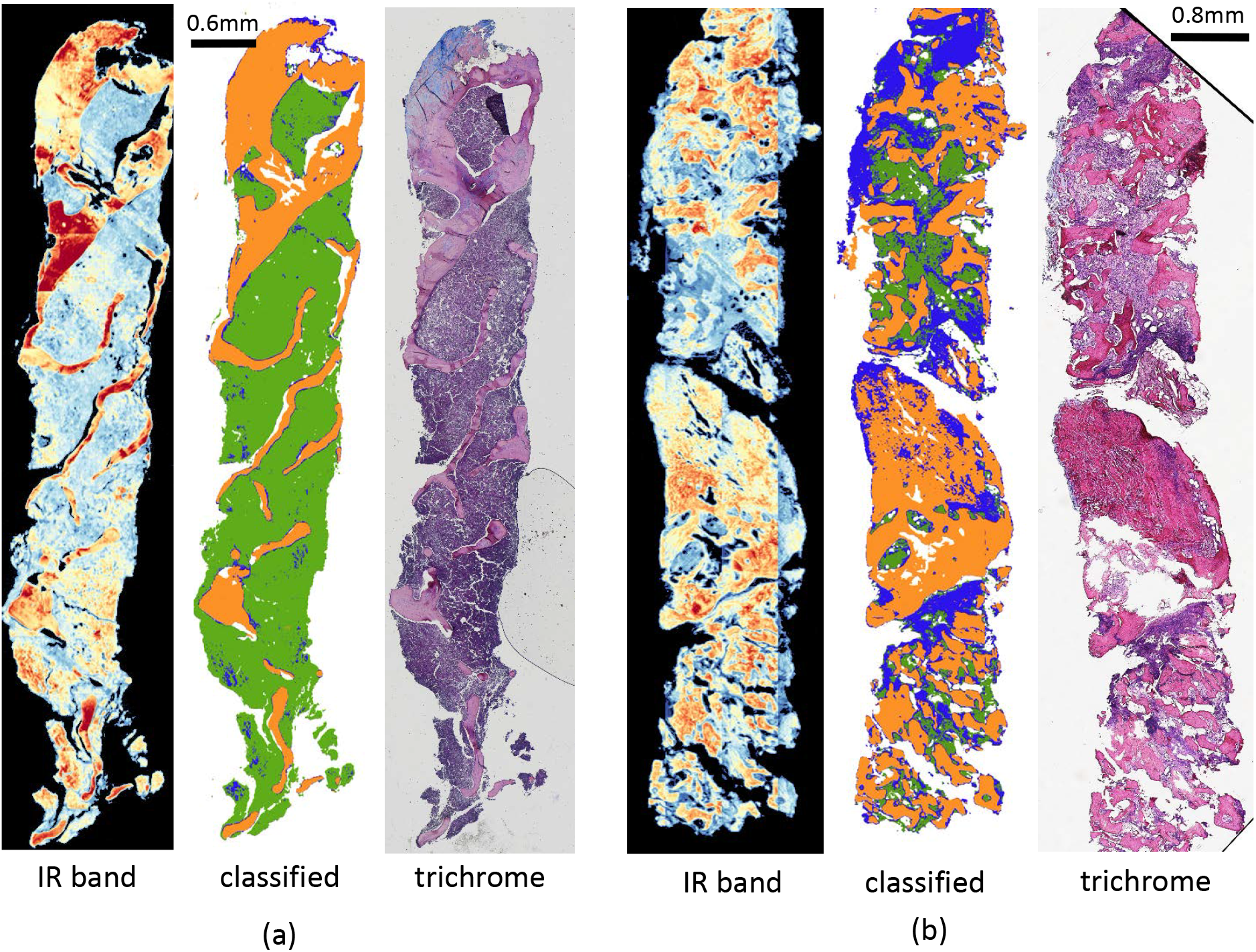
Classification results of FTIR imaged data for type I collagen (blue), trabecular bone (orange) and hematopoietic (green) and corresponding Mason’s trichrome stained adjacent tissue section for each grade.

We score collagen deposition by calculating the percentage of collagen pixels and the size of the largest cluster. Box plots for both metrics (Figure 12) show the separability of grades 0 and 1 from grades 2 and 3. Integrating both metrics allows differentiation of grades 0 and 1 from grades 2 and 3 with 89.6% accuracy. However, the accuracy for classification for all four grades reduces to 50% since differentiating grade 0 from grade 1 and grade 2 from grade 3 requires greater spatial resolution than what is available with diffraction-limited FTIR images.

**Figure 12:**
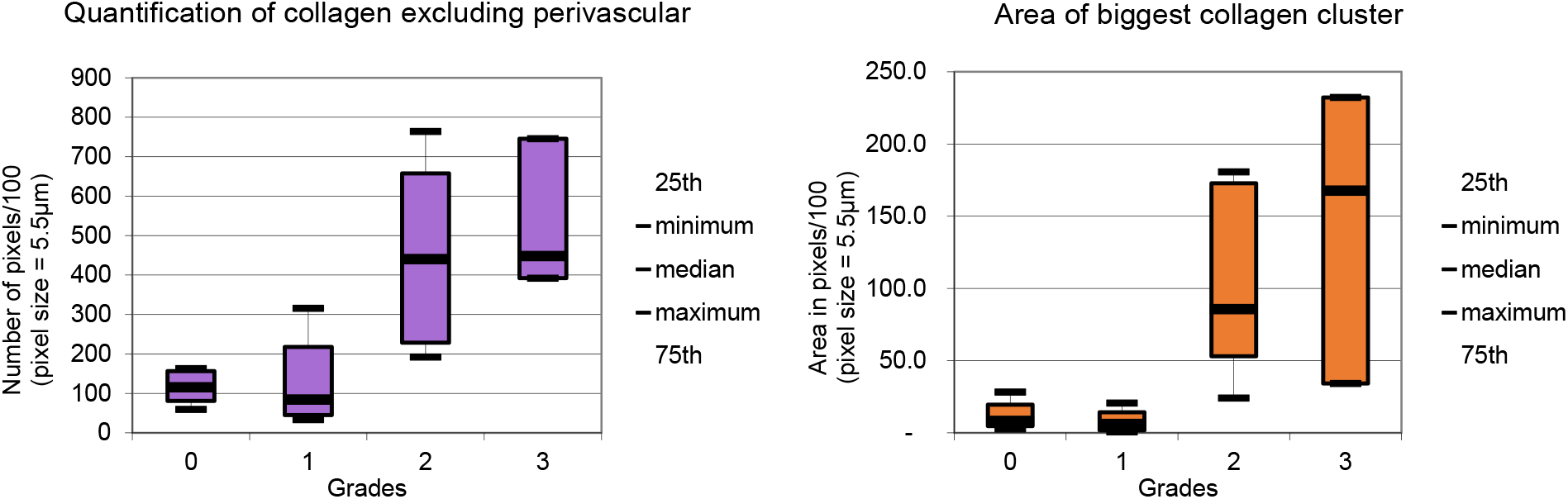
Box plots for spatial features quantifying collagen deposition in bone marrow biopsies.

## Conclusion

This work shows the potential for quantitative bone biopsy grading using infrared spectroscopic imaging. Osteosclerosis grading can be performed on standard glass slides, providing the potential for integration into existing pathology pipelines. The most optimistic approach for clinical integration is likely R-compatible substrates combined with DFIR imaging.^35^

Concurrence between pathologists is between 89.4%–94.9% for osteosclerosis grading and between 84.6%–91.3% for collagen deposition grading.^22^ The proposed method achieves accuracies of 84.4% for osteosclerosis grading using 19 samples and 88.88% using 18 samples (by eliminating traditionally-excluded artifacts). The accuracy for collagen deposition grading is 50%, which increases to 89.6% when considering only grades 0/1 and 2/3.

Spatial resolution introduces a critical limitation of infrared spectroscopic imaging for histological applications. While this is somewhat offset molecular specificity, many applications critically rely on spatial features. In particular, automated reticulin grading will likely require spatial resolution comparable to standard optical microscopes, which can potentially be achieved using O-PTIR imaging systems.^20,36^

## Acknowledgement

This work was funded in part by the Cancer Prevention and Research Institute of Texas (CPRIT) #RR140013, the National Heart, Lung, and Blood Institute #R01HL146745, and the National Library of Medicine #T15LM007093.

## References

(1) Thiele, J.; Kvasnicka, H. M.; Facchetti, F.; Franco, V.; van der Walt, J.; Orazi, A. Haematologica 2005, 90, 1128–1132.

(2) Swerdlow, S.; Campo, E.; Harris, N.; Jaffe, E.; Pileri, S.; Stein, H.; Thiele, J.; Arber, D.; Hasserjian, R.; Le Beau, M.; Orazi, A.; Siebert, R. WHO Classification of Tumours of Haematopoietic and Lymphoid Tissues. (revised 4th edition).; IARC, 2017; Vol. 2.

(3) Teman, C. J.; Wilson, A. R.; Perkins, S. L.; Hickman, K.; Prchal, J. T.; Salama, M. E. Leukemia Research 2010, 34, 871–876.

(4) Hipp, J.; Flotte, T.; Monaco, J.; Cheng, J.; Madabhushi, A.; Yagi, Y.; Rodriguez Canales, J.; Emmert-Buck, M.; Dugan, C. Michael; Hewitt, S.; Toner, T.; Tompkins, R.; Lucas, D.; Gilbertson, J.; Balis, U. Journal of Pathology Informatics 2011, 2.

(5) Veta, M.; Pluim, J. P.; Van Diest, P. J.; Viergever, M. A. IEEE Transactions on Biomedical Engineering 2014, 61, 1400–1411.

(6) Snead, D. R. J. et al. Histopathology 2016, 68, 1063–1072.

(7) Webster, J. D.; Dunstan, R. W. Veterinary Pathology 2014, 51, 211–223.

(8) Hoehn, D.; Medeiros, L. J.; Kantarjian, H. M.; Cortes, J. E.; Wang, X.; Bueso-Ramos, C. E. Human Pathology 2012, 43, 2354–2359.

(9) Berman, E.; Nicolaides, M.; Maki, R. G.; Fleisher, M.; Chanel, S.; Scheu, K.; Wilson, B.-A.; Heller, G.; Sauter, N. P. New England Journal of Medicine 2006, 354, 2006–2013.

(10) Old, O.; Lloyd, G.; Nallala, J.; Isabelle, M.; Almond, L.; Shepherd, N.; Kendall, C.; Shore, A.; Barr, H.; Stone, N. Analyst 2017, 142, 1227–1234.

(11) Mayerich, D. M.; Walsh, M.; Kadjacsy-Balla, A.; Mittal, S.; Bhargava, R. Breast histopathology using random decision forests-based classification of infrared spectroscopic imaging data. Proc. SPIE–Int. Soc. Opt. Eng. 2014; p 904107.

(12) Benard, A.; Desmedt, C.; Smolina, M.; Szternfeld, P.; Verdonck, M.; Rouas, G.; Kheddoumi, N.; Rothé, F.; Larsimont, D.; Sotiriou, C.; Goormaghtigh, E. Analyst 2014, 139, 1044–1056.

(13) Baker, M. J.; Gazi, E.; Brown, M. D.; Shanks, J. H.; Gardner, P.; Clarke, N. W. British Journal of Cancer 2008, 99, 1859–1866.

(14) Bassan, P.; Sachdeva, A.; Shanks, J. H.; Brown, M. D.; Clarke, N. W.; Gardner, P. Automated high-throughput assessment of prostate biopsy tissue using infrared spectroscopic chemical imaging. Proc SPIE. 2014; pp 90410D–90416D.

(15) Großerueschkamp, F.; Kallenbach-Thieltges, A.; Behrens, T.; Brüning, T.; Altmayer, M.; Stamatis, G.; Theegarten, D.; Gerwert, K. Analyst 2015, 140, 2114–2120.

(16) Ozek, N. S.; Tuna, S.; Erson-Bensan, A. E.; Severcan, F. Analyst 2010, 135, 3094–3102.

(17) Belbachir, K.; Noreen, R.; Gouspillou, G.; Petibois, C. Analytical and Bioanalytical Chemistry 2009, 395, 829–837.

(18) Hammiche, A.; Pollock, H.; Reading, M.; Claybourn, M.; Turner, P.; Jewkes, K. Applied Spectroscopy 1999, 53, 810–815.

(19) Li, Z.; Aleshire, K.; Kuno, M.; Hartland, G. V. The Journal of Physical Chemistry B 2017, 121, 8838–8846.

(20) Reffner, J. A. Spectroscopy 2018, 33, 12–17.

(21) Wilkins, B. S.; Erber, W. N.; Bareford, D.; Buck, G.; Wheatley, K.; East, C. L.; Paul, B.; Harrison, C. N.; Green, A. R.; Campbell, P. J. Blood 2008, 111, 60–70.

(22) Kvasnicka, H. M.; Beham-Schmid, C.; Bob, R.; Dirnhofer, S.; Hussein, K.; Kreipe, H.; Kremer, M.; Schmitt-Graeff, A.; Schwarz, S.; Thiele, J. Histopathology 2016, 68, 905–915.

(23) Kuter, D. J.; Bain, B.; Mufti, G.; Bagg, A.; Hasserjian, R. P. British Journal of Haematology 2007, 139, 351–362.

(24) Yagi, Y. Color standardization and optimization in whole slide imaging. Diagnostic Pathology. 2011; p S15.

(25) Walsh, M. J.; Mayerich, D.; Kajdacsy-Balla, A.; Bhargava, R. High-resolution mid-infrared imaging for disease diagnosis. Biomedical Vibrational Spectroscopy V: Advances in Research and Industry. 2012; p 82190R.

(26) Tiwari, S.; Bhargava, R. The Yale Journal of Biology and Medicine 2015, 88, 131–143.

(27) Frost, J.; Ludeman, L.; Hillaby, K.; Gornall, R.; Lloyd, G.; Kendall, C.; Shore, A. C.; Stone, N. Analytical Methods 2016, 8, 8452–8460.

(28) Wrobel, T. P.; Bhargava, R. Analytical Chemistry 2017, 90, 1444–1463.

(29) Mankar, R.; Walsh, M.; Bhargava, R.; Prasad, S.; Mayerich, D. Analyst 2018, 143, 1147–1156.

(30) Bassan, P.; Mellor, J.; Shapiro, J.; Williams, K. J.; Lisanti, M. P.; Gardner, P. Analytical Chemistry 2014, 86, 1648–1653.

(31) Fernandez, D. C.; Bhargava, R.; Hewitt, S. M.; Levin, I. W. Nature Biotechnology 2005, 23, 469–474.

(32) Baker, M. J. et al. Nature Protocols 2014, 9, 1771–1791.

(33) Pilling, M. J.; Henderson, A.; Shanks, J. H.; Brown, M. D.; Clarke, N. W.; Gardner, P. Analyst 2017, 142, 1258–1268.

(34) Baker, M. J.; Byrne, H. J.; Chalmers, J.; Gardner, P.; Goodacre, R.; Henderson, A.; Kazarian, S. G.; Martin, F. L.; Moger, J.; Stone, N. Analyst 2018, 143, 1735–1757.

(35) Kuepper, C.; Kallenbach-Thieltges, A.; Juette, H.; Tannapfel, A.; Großerueschkamp, F.; Gerwert, K. Scientific Reports 2018, 8, 7717.

(36) Zhang, D.; Bai, Y.; Cheng, J. Photonics Media 2018,

